# Robust frequency-encoded dynamics in a minimal synthetic phytohormone crosstalk

**DOI:** 10.1101/2020.05.31.125997

**Authors:** S. Pérez García, M. García Navarrete, D. Ruiz Sanchis, C. Prieto Navarro, M. Avdovic, O. Pucciariello, K. Wabnik

**Affiliations:** Centro de Biotecnologıa y Genomica de Plantas (Universidad Politecnica de Madrid – Instituto Nacional de Investigacion y Tecnologıa Agraria y Alimentaria), Autopista M-40, Km 38 – 28223 Pozuelo de Alarcon, Spain

## Abstract

How do dynamic hormone inputs translate into speed, and precision of response is one of the most challenging questions of science. To approach this question, we constructed minimal synthetic gene circuits capable of responding to plant hormones auxin and salicylic acid (SA). These circuits integrate bacterial multi antibiotic resistance (Mar) repressors that directly detect phytohormones through a ligand-induced conformational switch. The combination of individual circuits in synthetic auxin-SA crosstalk was sufficient to coordinate responses across the cell population with tunable precision and speed in long-term microfluidics experiments. This antagonistic auxin-SA crosstalk retains temporal memory upon extended exposure to hormones and synchronizes the behavior of individual cells with the environmental clock. Our study shows how dynamic hormone inputs can be translated in robust and precise responses with a minimal assembly of bacterial transcriptional repressors, suggesting an alternative regulatory strategy to known plant hormone signaling systems.

## Introduction

Auxin and salicylic acid (SA) are small signaling molecules critical for growth and immune responses in plants **[1-3]**. Natural biosensors for auxin an SA have been used to understand mechanisms of complex plant hormone action **[4-8]**. Nevertheless, such systems are indirect **[9]** and depend on several plant-specific components **[4,5]** or entire microbes **[6]**. For example, auxin biosensors have been primarily developed for targeted protein degradation in plant and non-plant systems **[8, 10, 11]**. Neither of these systems, however, offer a tunable regulation of input-output relation under a dynamically changing environment. Minimal genetic systems for direct sensing of auxin and SA would offer opportunities for creating new regulatory modules that translate dynamic hormone inputs into programmable outputs. To implement and test such systems, baker’s yeast *S. cerevisiae* provides an ideal host because it does not utilize auxin nor SA through dedicated signaling networks.

Potential elements for engineering minimal systems of phytohormone signaling could be identified in plant-associated microbes. This is because, many symbiotic or pathogenic microbes have the capacity of metabolizing phenol and indole-based compounds through multiple antibiotic resistance (Mar) regulons **[12]**. *E.coli* MarR transcriptional repressor is a representative member of this family (Supplementary Fig. 1) that is required for antibiotic stress response in the presence of salicylates **[13, 14]**. In contrast, the MarR ortholog from *A. Baumanii* (IacR) is sensitive to naturally occurring auxin such as 3-indoleacetic acid (IAA) **[15]**. It has been proposed that SA and IAA control the activity of MarR and IacR repressors **[12-15]**, however, the mechanistic basis of such sensing remains unclear.

Guided by insights in phytohormone-dependent regulation of Mar proteins provided by molecular dynamics simulations, we constructed genetic circuits for direct sensing of auxin and SA using a pair of orthogonal MarR transcriptional repressors. These minimal synthetic circuits record changes in physiological concentrations of both hormones in individual cells with dynamic range, specificity and orthogonality. To probe the input-output relation in a quantitative matter we constructed the synthetic crosstalk of auxin-SA by coupling circuits to a downstream output. This auxin-SA crosstalk generated tunable and synchronous responses to dynamic hormone changes in the precisely controlled environment on the microfluidic chip. Our simple phytohormone crosstalk system is capable of ‘programming’ complex dynamics including logic gating and oscillatory response tuned to an external clock, providing compact yet powerful alternative to natural plant hormone signaling networks.

## Results and Discussion

### MarR repressors undergo structural transitions from flexible to rigid conformation in the presence of hormones

We initially address the elusive dynamics of ligand sensing by MarR repressors with all-atom molecular dynamics (MD) simulations performed on MarR crystal structures. MD simulations demonstrate that MarR apoprotein remained flexible (Fig. 1a,d) and adapt to the shape of the DNA operator by adjusting a distance between DNA binding (DB) domains (Fig. 1b, Supplementary Movies 1 and 2). On the contrary, the presence of SA in the MarR dimerization interface brought monomers close to each other, creating a rigid closed structure (Fig. 1c,d, Supplementary Movie 3) that was unable to bind DNA. We then calculated the probability distribution of the distance between R73 residues of DB domains that are in direct contact with DNA. We found that only apoprotein, largely due to its flexibility, was able to scan across various distances and eventually sat at the DNA-bound state (Fig. 1e). To address the dynamics of conformation changes, we performed the metadynamics energy landscape analysis with the local perturbation of distance between R73 atoms of both monomers (see Materials and Methods for details).

**Figure 1.**
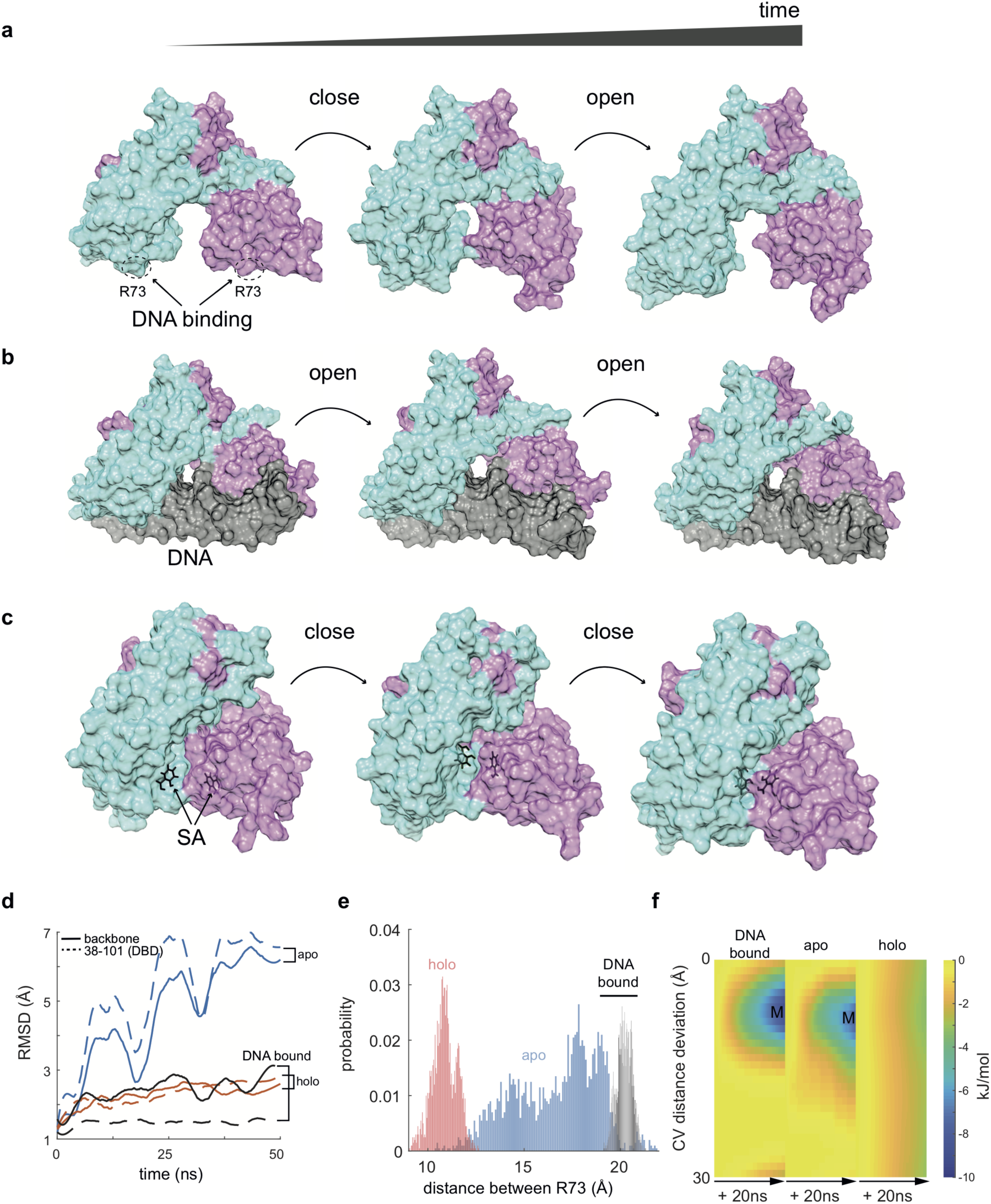
A conformational switch drives the phytohormone sensing through MarR-type regulators. (**a-c**), Time-lapse from Molecular Dynamics Simulations of the flexible MarR apoprotein (monomers are shown in cyan and purple) (a) (RCSB: 5h3r and 3voe) display a cyclic change of conformation states (open-closed-open). MarR dimer bound to DNA operator site (black) remains in the open conformation with a specific distance of ∼ 20 A ° between DNA contacting R73 (b). MarR bound to two molecules of SA (c) (black, RCSB: 1jgs) display successive shortening of the distance between R73 that reflects an unbinding of MarR from DNA. (**d**), Time-trace of molecular dynamics simulations with MarR_apo_ (blue), MarR_holo_ (red) and DNA-bound MarR (black). The root-mean-square deviation of atomic positions (RMSD) is shown for dimer simulations of 50 ns. Note, the cyclic pattern of closed-to-open conformation of MarR_apo_ which was absent from MarR_holo_ simulations as holoprotein stabilizes in the closed conformation. In contrast, DNA-bound MarR retains an open confirmation. (**e**), A probability distribution of the mean distance between R73 atoms of both monomers for apo (blue), holo(red), and DNA-bound (black) configurations. Note that the holo state provides a ≈ 2-fold short distance between DBD domains than that required for a stable bond with DNA. (**f**), Energy landscapes for three MarR configurations (apo, holo, DNA-bound) for deviation from the preferred distance of ≈ 20 A ° between R73 (both chains) required for DNA binding. Note energy minima wells (M) for apo and DNA bound configurations which were absent in holoprotein simulations indicating that the hormone-bound MarR did not reach an energy minimum required for the association with the DNA.

Time-dependent changes of energy portraits in apoprotein and DNA-bound MarR configurations revealed the presence of a stable energy minimum (M), which was absent in holoprotein simulations (Fig. 1f). This minimum corresponds to the smallest deviation from the reference distance of ≈ 20 A° between DB domains, as observed in crystals of DNA-bound MarR (Fig. 1e). Collectively, our findings indicate that MarR repressor requires high flexibility to adapt to DNA operator sites which is negatively regulated by phytohormone exposure.

Next, we asked if IacR could respond to auxin using a similar mechanism to that of MarR. The model of IacR predicted that apoprotein remains flexible concerning the spacing between DB domains (Supplementary Fig. 2a). In contrast, the bound IAA stretched the distance between these domains (Supplementary Fig. 2b,c), as demonstrated by the frequency of distance distribution (Supplementary Fig. 2d) and metadynamics analysis (Supplementary Fig. 2e). Therefore predicted IacR dynamics share common design principles of DNA recognition with MarR, including backbone flexibility and successive switches of conformation in the presence of hormone.

### Synthetic circuits robustly respond to temporal graded changes of auxin and SA

We tested whether MarR and IacR would be suitable candidates for building minimal synthetic phytohormone signaling systems in the eukaryotic host. MarR or IacR were introduced in yeast, driven by galactose induction with a strong herpes simplex virus trans-activation domain (VP64) **[16]** and PEST degron **[17]** for fast degradation dynamics (Fig. 2a and Supplementary Fig. 3a). We used a unstable Green fluorescent reporter (dEGFP) **[18]** expressed from the synthetic minimal promoter with putative MarR **[13]** or IacR operator sequences **[15]**. Notably, both IacR and MarR showed a dynamic range of response (up to 2-orders of magnitude) to physiological phytohormone concentrations (Supplementary Fig. 3c,d) with EC_50_ = 0.13 μM (IAA) and 407.3μM (SA), respectively. Furthermore, IacR module response was highly specific to IAA and insensitive to SA (Supplementary Fig. 3e) whereas MarR module did not respond to IAA (Supplementary Fig. 3f), confirming the orthogonality of both phytohormone sensing circuits.

**Figure 2.**
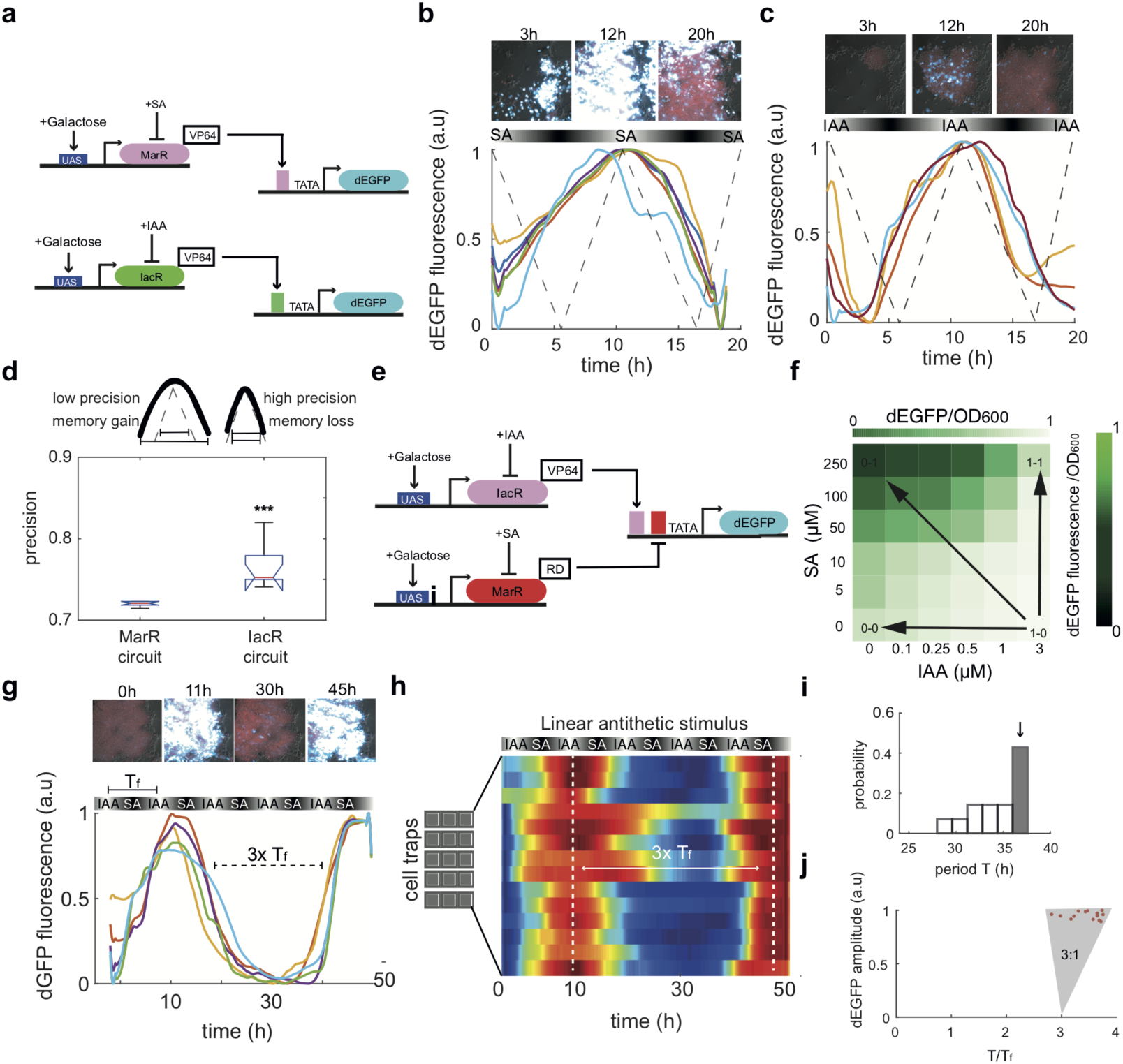
Synthetic MarR-based gene circuits integrate physiological concentrations of phytohormones in a dynamically changing environment. (**a**), Schematics of SA (top) and IAA (bottom) sensing circuits. MarR or IacR repressors are tagged with a synthetic VP64 viral transactivation domain, nuclear localization signal (NLS), and PEST degron. Galactose is used to induce the transcription of MarRs. The fluorescent reporter (dEGFP) contains an uncleavable ubiquitin (G76V) tagged to N-terminus of enhanced GFP (EGFP) that is driven by the minimal promoter with either MarR(pink bar) or IacR(green bar) operators placed directly upstream of TATA-box. (**b, c**), Examples of screenshots and normalized dEGFP (cyan) time traces for SA (b) and IAA circuits (c) across different traps from independent microfluidic experiments integrating linear changes of phytohormones (dashed lines). Cells carrying MarR-driven circuit (b) displayed a broader peak and delayed response to a cyclic linear change in hormone concentrations (10 hours period of SA or IAA) compared to cells equipped with the synthetic auxin sensor (c). (**d**), Circuit precision is inversely correlated with a temporal memory or ‘capacitance’ of the previous hormone state. Comparison of precision between IAA and SA circuits calculated for at least n**=**10 independent traps per circuit tested using a one-way ANOVA test (p*** <0.001). (**e**), Implementation of the CrossTalker system. MarR was tagged with the repression domain consisting of the last 24 amino acids of MIG1 transcriptional repressor, nuclear localization signal (NLS), and mouse ODC degron and combined with activating IacR-VP64-ODC construct. A synthetic promoter that drives dEGFP, consists of IacR and MarR operators upstream of TATA-box. **(f)**, Dual gradient scans on CrossTalker reveals four boundary states (0-0; 1-0;0-1; 1-1) reminiscent of the NIMPLY logic gate (average of triplicates is shown). Color coding map for normalized dEGFP fluorescence is shown. (**g, h**), Examples of CrossTalker circuit dynamics in yeast cells (g) that were grown in the microfluidic device with 10-hour antithetic waves of SA and IAA. Note, a consistent synchronization across 14 independent trapping regions (h). (**i**), A histogram of mean period distribution from 14 traps on the microfluidics chip. Gray bar shows a dominant frequency of response. (**j**), The plot of mean dEGFP fluorescence amplitude against the ratio of response (T) and input forcing (T_f_) periods in 14 independent traps. The shaded synchronization region; 3 periods of stimulus corresponds to a single period of CrossTalker circuit response.

Next, we probed the dynamics of phytohormone-driven circuits in a dynamic environment that is governed by non-static changes in hormone levels. For that purpose, we developed bench-top customized rapid, cost-effective lithography technologies **[19]** for the fabrication of microfluidic devices (Supplementary Fig. 4a). Our microfluidics devices (Supplementary Fig. 4a) integrates a time-dependent switching of two independent media inputs, through a chaotic mixer module (Supplementary Fig. 4b,c). Next, we performed a chaotic microfluidic mixing with a linear decrease and subsequent increase of SA and IAA in 10h cycles (Fig. 2b,c, dashed lines). We could observe, in our microfluidic experiments, a prominent pulse of reporter gene dEGFP synchronized across the large population of exponentially growing cells (∼1000 cells per trap of size 500μm x 500μm) (Fig. 2b,c and Supplementary Fig. 5a-c). We then looked into the precision of MarR and IacR circuits which was defined as relative width of response peak compared to width of hormone input pulse (Fig. 2d). Our findings indicate the trade-off between memory and precision of responses exists such that an increased precision of response involves the loss of temporal memory of previous phytohormone concentrations. We found that both circuits showed ∼72% precision, however, the IacR module showed ∼5% higher precision than that of MarR (Fig. 2d).

### Synthetic phytohormone crosstalk adapts robustly to dynamic changes in the environment

Since both Mar-based circuits are orthogonal, we attempted to combine them to create synthetic crosstalk of phytohormones to which we refer as ‘CrossTalker’ (Fig. 2e). The general scheme of CrossTalker architecture includes the IacR-VP64 activator and MarR repressor directly fused to the minimal repression domain from yeast MIG1 transcriptional regulator **[20]** (Fig. 2e and Supplementary Fig. 3b). The CrossTalker displayed a robust NIMPLY gated response to step-wise elevations of both phytohormones in microwell plate fluorescence scanning assays (Fig. 2f), confirming the orthogonality of both sensor modules. Similarly, a gated response was observed in the microfluidic device exposed to combinatorial inputs (Supplementary Fig. 5d).

To further probe Crosstalker circuit dynamics with our microfluidic platform we performed hormone mixing over the time course of the experiment with a double linear gradient generator shifted in phase by π with a period of 10 hours. The CrossTalker circuit (Fig. 2e) exhibited dynamic waves of reporter gene expression (Fig. 2g, h, Supplementary Movie 4). Reporter peaks were separated by approximately three periods of an input signal (Fig. 2i,j) and showed synchronization across traps (Fig. 2h,j) with the negligible variation in peak amplitudes (Supplementary Fig. 5e). To understand the cause of a delay between consecutive response peaks observed in our experiments we constructed a computer model of hormone crosstalk (Supplementary Fig. 5f-h). Model predicted that a delay in the repression pathway (MarR-RD) could produce a similar output (Supplementary Fig. 5h) to that observed experimentally (Fig. 2g).

### Increased frequency of environmental clock fine-tunes precision and speed of a synthetic auxin-SA crosstalk

Finally, we tested whether increasing the frequency of stimulus could potentially lead to increased speed and precision of CrossTalker responsiveness, providing the frequency entrainment mechanism (Fig. 3a). Our computer model simulations of the CrossTalker circuit predict a correct synchronization with the cyclic phytohormone stimulus (Supplementary Fig. 5i). To test these predictions experimentally the Crosstalker circuit was subjected to 5-hour antithetic pulses of IAA and SA which constitute the environmental clock (Fig. 3b). On-chip imaging revealed robust, synchronized oscillations of a reporter gene (Fig. 3c, Supplementary Movie 5) with a mean period of 5-6 hours (Fig. 3d) and minimal variation of the amplitude (Supplementary Fig. 6a). Similarly, the autocorrelation analysis confirmed synchronized oscillations within a narrow frequency domain (Fig. 3e) with two synchronization modes 1:1 and 1.5:1 (Supplementary Fig. 6b). Also, microfluidics experiments revealed a phase diffusion of the period that may impact the long-term precision of circuit response (Supplementary Fig. 6c). The successive increase of hormone input frequency by 2-fold led to faster, sharper and more consistent oscillations (Fig. 3f, Supplementary Movie 6) that were synchronized across different microfluidics traps (Fig. 3g). Consistently, the autocorrelation of reporter signal revealed the prevalence of oscillations with shorten periods (between 3 and 5 h) (Fig. 3h,i and Supplementary Fig. 6d) compared to those observed with 2x slower stimulus (Fig. 3d). No significant delay between successive periods was observed (Supplementary Fig. 6e). By inspecting in-depth the response characteristics of the Crosstalker circuit, we found that the mean dEGFP amplitude per trap was reduced by 25% (Supplementary Fig. 6f) while the consistency of period was significantly improved (Supplementary Fig. 6g) towards the higher frequency of the hormone input. Finally, the precision of CrossTalker was increased by 15% (Supplementary Fig. 6h) whereas the Coefficient of variation (CV) dropped by almost 2-fold (Supplementary Fig. 6i). Therefore, faster changes in the hormone inputs improve the speed and precision of response at the cost of temporal memory loss (Fig. 3j).

**Figure 3.**
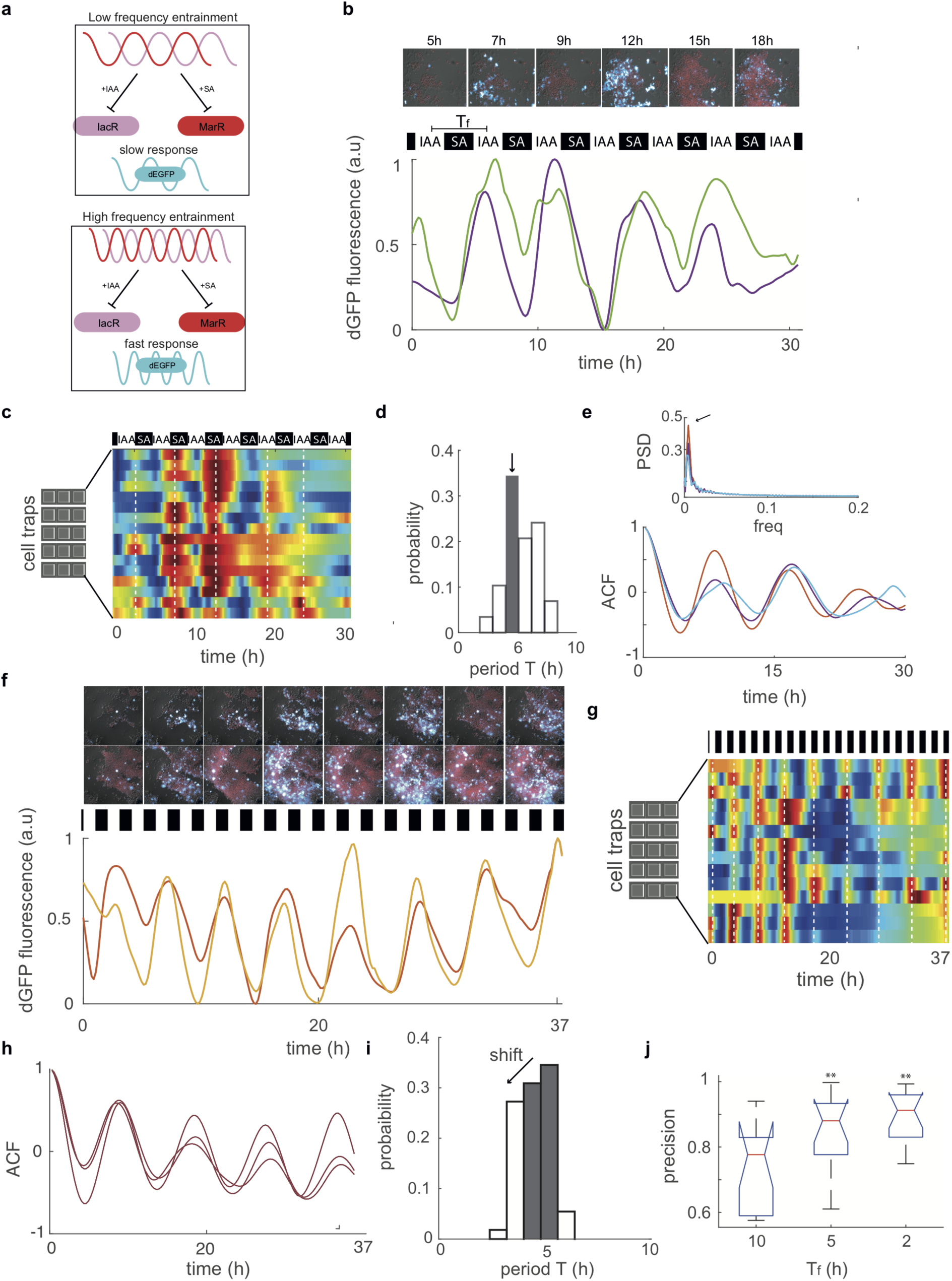
CrossTalker circuit synchronizes the response of cell populations to the environment clock through the precision frequency-encoded mechanism. (**a**), A cartoon demonstrates the principles of frequency entrainment mechanism for the Crosstalker circuit. For slow cycles of the input stimulus, a circuit displays a slow response (upper panel). The increased frequency of the stimulus is decoded by the circuit to adjust the speed of response (lower panel). (**b, c**), Dynamics of crosstalk circuit response towards the 5h antithetic hormone pulses (b) show a fair synchronization across traps n=16 traps (c), revealing input frequency-dependent tunability of the CrossTalker circuit. (**d**), A period probability distribution displays a dominant frequency (gray bar) of response locked to the input signal frequency. (**e**), Autocorrelation (ACF) (lower panel) and power spectrum density (PSD) traces(top panel) across different traps demonstrate a periodic character of circuit response with a dominant frequency peak (top panel). (**f**), Image snapshots (every 2h), and time traces of circuit oscillations with rapid 2-hour antithetic hormone pulses. (**g**), Improved synchrony across cell populations shown in n=14 traps in the microfluidic chip. (**h**), cumulative ACF plots for example traps show a sustained periodic pattern. (**i**), A period probability distribution displays a shift to shorter period or higher frequency (gray bars, arrow). (**j**), Comparison of circuit precision for three frequencies of hormone stimuli. Precision and speed of response increased with the higher frequency of applied hormone inputs, causing the memory loss (compare to Fig. 2g). Comparison of precision between Crosstalker circuit in dynamic environments calculated for at least n**=**14 independent traps using a one-way ANOVA test (**p < 0.01).

Taken together, our data demonstrate that our CrossTalker system converts dynamic fluctuations in the environment into a tunable response by decoding the dominant frequency of the hormone stimulus. This minimal synthetic crosstalk generates either temporal memory in hormone concentrations or response with increased precision and speed that depend on the frequency of environmental changes.

A mechanistic mapping of dynamic environmental inputs into precise outputs is a key to both understand and engineer phenotypic patterns in biology. In plants, phytohormones auxin and SA play pivotal roles in the regulation of growth and immunity **[21-26]**, offering plant resilience to different types of environment. However, plant signaling networks involving auxin and SA, are complex and demonstrate a sophisticated multiple layered organization. To provide a simpler alternative to these natural networks, we engineered minimal synthetic circuits for direct sensing of auxin and SA from MarR-type repressors from bacteria **[12]**. The combination of structural insights, computer modeling, and microfluidics revealed that circuits robustly sense graded concentration of hormones with high temporal resolution. Our work lays mechanistic grounds for the quantitative assessment of transient phytohormone dynamics and engineering of gene circuits that are capable of decoding spatial-temporal hormone information into precise response times. Proposed phytohormone-driven circuits can be directly ported to any eukaryotic or prokaryotic host, expanding the range of applications for engineering more complex regulatory systems.

## Acknowledgments

We are grateful to L. Rubio for sharing yeast strain CEN.PK2-1C. Authors thank Mark Estelle for critical discussions at the early stage of the project. This work was supported by the Programa de Atraccion de Talento 2017 (Comunidad de Madrid, 2017-T1/BIO-5654 to K.W.), Severo Ochoa (SO) Programme for Centres of Excellence in R&D from the Agencia Estatal de Investigacion of Spain (grant SEV-2016-0672 (2017-2021) to K.W. via the CBGP). In the frame of SEV-2016-0672 funding M.A. received a PhD fellowship (SEV-2016-0672-18-3: PRE2018-084946) and O.P. is supported with a postdoctoral contract. K.W. was supported by Programa Estatal de Generacion del Conocimiento y Fortalecimiento Cientifico y Tecnologico del Sistema de I+D+I 2019 (PGC2018-093387-A-I00) from MICIU (to K.W.).

## Author contributions

S.P.G and M.G.N designed, performed most of experiments and analyzed data. D.R.S, C.P.N and O.P contributed to plasmid and strain constructions, S.P.G and M.A, performed multi-well plate fluorescence assays. M.G.N performed microfluidics experiments. K.W designed experiments analyzed data and supervised the project. All authors contribute to the manuscript writing.

## Data and materials availability

All data that support the findings of this study are available in the manuscript. Original data that supports the findings are available from the corresponding author upon request. Plasmids sequences will be deposited in NCBI database after publication. Correspondence and requests for materials such as should be addressed to K.W. Image processing and analysis were performed using Fiji (v.1.0) and Matlab (R2018b) and R-studio. Subsequent analysis was performed using custom R and Matlab scripts. The codes are available upon request from the corresponding author.

## Materials and Methods

### Structural modeling and Molecular dynamics simulations

Crystal structures of the apoprotein, DNA-bound and holoprotein were obtained from Protein Data Bank RCSB database (codes: 5h3r, 3voe, and 1jgs). In preparation for MD simulations, structures were process using Charmm-gui (www.charmm-gui.org) Solution Builder. Simulation-ready structures contain hydrogens, water box around the structure fitted to the size of system, and water environment with 0.15M NaCl ions and an isothermal-isobaric (NPT) ensemble with a constant amount of substance (N), pressure (P) and temperature (T). The pressure was set at 1 atm (1.01325 bars) and temperature at 303 K. MD simulations were performed using OpenMM environment (http://openmm.org/) with hydrogen mass repartitioning (HMR) scheme. Initial steps of minimalization and equilibration for 1 ns were followed by tracking 50 ns conformation dynamics with the time step of 2 femtoseconds. Particle Mesh Ewald (PME) periodic boundary conditions were used for the electrostatic force calculations. Trajectory files from the simulations were processed with VMD (www.ks.uiuc.edu/Research/vmd/) to generate RMSD plots and atom distance distributions. The visualization of structures was done in UCSF Chimera (https://www.cgl.ucsf.edu/chimera/). Metadynamics energy landscape analysis was performed in OpenMM using Metadynamics module and Morse/long-distance potential between atomic centroids of R73 (MarR) and K80 (iacR) residues in DB domains. This collective variable (CV) is calculated based on the reference distance of ≈ 20 A° inferred from the crystal structure of DNA-bound MarR as en equilibrium parameter in the Morse potential function. 50ns trajectories were used as a starting point to perform an additional 20ns simulation (total 70ns) with CV in the flexible range of 0-3 nm. The IacR structure models were obtained with Swiss-Model using several templates, RCSB codes: 1jgs, 5hr3 and 3voe. The molecular docking of IacR with two IAA molecules was performed using UCSF Chimera and the AutoDock Vina. IacR and IAA charges were merged and the non-polar hydrogens and lone pairs were removed. In addition, hydrogens were added to the IacR structure. A total of 10 binding modes were generated and best two poses were selected based on the minimal energy score. The exhaustiveness of the search was set to 8.

### Structure alignment

Molecular Evolutionary Genetics Analysis (MEGA) was used to produce the alignment of the multiple MarR proteins, specifically the ClustalW program available in MEGA. The Gap Opening Penalty was set at 10.00 and the Gap Extension Penalty at 0.20. The protein weight Matrix used in this multiple alignments was Gonnet. For the visualization and generation of the alignment image, we used the Jalview program with the input alignment file from MEGA.

### Strains and plasmid constructions

All plasmids in this study were created using isothermal Gibson assembly cloning **[27]**. A middle-copy (15-30 copies) episomal plasmid pGADT7 (Takara Bio) was used to increase the concentration of synthetic circuit components in order to buffer for the intrinsic molecular noise. This plasmid was further modified to accommodate three different auxotrophic selection markers (Leucine, Uracil, and Histidine; Supplementary Fig. 3a,b). GAL7 promoter and either CYC1 or ADH1 terminators were cloned and combined with MarR modules in activator or repressor plasmids (Supplementary Fig. 3a,b). The reporter plasmids include synthetic minimal promoters with putative MarR or IacR operator sequence upstream TATA-box and fast-degradable UBG76V-EGFP (dEGFP). PCR reactions were performed using Q5 high fidelity polymerase (New England Biolabs). Correct PCR products were digested with DpnI (New England Biolabs) to remove the template and subsequently cleaned up with a DNA cleanup kit (Zymo Research) before Gibson assembly. Constructs were transformed in ultra-competent cells from E. coli DH5a strain using standard protocols. All plasmids were confirmed by colony PCR and validated with sequencing. The CEN.PK2-1C (a kind gift from Dr. Luis Rubio) yeast strain was used to make competent cells and transformation of plasmids using Frozen-EZ Yeast Transformation II Kit (Zymo Research).

### Multiwell plate fluorescence measurements

Overnight culture of the yeast in 2% sucrose low fluorescence media (Formedium, UK) was diluted 100 times and transfer to a 96 well plate with sucrose only or 2% sucrose + 0.5% galactose conditions and a 2D gradient of phytohormone concentrations. Plates were incubated at 30C overnight in the thermal incubator and well mixed by shaking before performing measurements. Measurements were done with the Thermo ScientificTM VarioskanTM LUX multimode microplate reader. Optical Density was set at an absorbance of 600nm wavelength, the fluorescence excitation and emission light at 488nm and 517nm wavelength, respectively.

### Microfluidic Mold fabrication

Molds for the production of microfluidic devices were designed in Adobe Illustrator software and printed on plastic sheets with the monochrome laser printer at 1200dpi resolution. A density of Ink deposition was used to control the feature height. Plastic wafers were cut and transferred to the thermal oven set to 160°C to shrink by one-third of the original size, the baked again for 10 min to smoothen and harden the ink. Finally, molds were cleaned with soap, rinsed with isopropanol and DDI water and dry using a nitrogen gun and secured with Scotch tape before use.

### Soft Lithography

Molds were introduced in plastic 90mm Petri dishes and fixed with double-sided tape. Dowsil Sylgard 184 Polydimethylsiloxane (PDMS) was properly mixed in a 10:1 (w/w) ratio of elastomer and curing agent and stirred until the uniform consistency was achieved. Approximately 27mL of the homogeneous mixture was poured in each petri dish and completely degassed using the 8 CFM 2-stage vacuum pump for approximately 20 minutes. Degassed PDMS was cured at 80°C for 2h. Cured PDMS was removed from the petri dish, separated from the wafer and cut to extract the individual chips. Fluid access ports were punched with 0.7mm diameter World Precision Instruments (WPI) biopsy puncher and flushed with ethanol to remove any remaining PDMS. Individual chips were cleaned with ethanol and DDI water and Scotch tape to remove any remaining dirt particles.

### Microfluidic device bonding

At least one day before use, individual chips and coverslips were cleaned in the sonic bath and rinsed in ethanol, isopropanol, and water. Both surfaces were exposed to Corona SB plasma treater (ElectroTechnics Model BD-20AC Hand-Held Laboratory Corona Treater) between 45 seconds to 1 minute, then surfaces were brought together and introduce at 80°C in an oven overnight to obtain the enhanced bond strength.

### Calibration of microfluidic mixer module

Mixer Calibration was performed by staining one of two inputs with Rhodamine B 0.001% (w/v). A pair of 60mL syringes were prepared with 20mL of water or water + dye. Three 50mL falcon tubes were filled with 20mL of water and connected to waste and cell loading ports. Syringes and falcon tubes were set on linear actuators which control pressure in the microfluidic device through customized software. The microfluidic device was vacuumed for at least 20 minutes to remove air bubbles and facilitate device wetting before connecting Tygon microbore tubing 0.020” x 0.060”OD (ColePalmer Inc.). After air removal, one syringe was connected and then the other syringes were plugged sequentially once liquid reached and filled the ports. Switching between inputs was regulated with the gravity-aided hydrostatic pressure by changing the relative height of syringes. Increasing height of input media 1 (M1) over input media 2 (M2) produces the pressure difference in the mixing module. Therefore, at maximum height differences, microfluidic chip is filled with 100% of M1 and 0% of M2. Syringe positions were adjusted with 0.1 mm precision by changing height to obtain highly efficient mixing between 0% and 100%. The excessive flow is diverted towards an auxiliary waste output (W1) that releases the pressure from the mixer. Flow then was introduced into the second chaotic mixer to assure a rapid switching of conditions. The calibration process was repeated at least three times with three independent microfluidic devices. Maximum and minimum height settings varied solely by a few millimeters, therefore we used averaged measures without cross-contaminating media inputs, which were following: 610 mm for the maximum and 390 mm for minimum settings, respectively. All three replicates showed exact 50% mixing for the 500 mm settings.

### Cell loading procedure

All tubing lines were sterilized with ethanol and plugged to syringes or introduced in falcon tubes under sterile conditions. Fresh yeast colony was grown in low fluorescence media composition (Formedium, UK) with 2% sucrose as a carbon source. The next day, yeast cultures were diluted 10-20 times approximately to 0.2-0.4 O.D_600_ to obtain highly concentrated cells that were transferred to a 50mL falcon tube for loading. 60mL media syringes were filled with 25mL of media with or without hormones and 50mL waste falcons tubes were filled with 10mL of DDI water. Before loading, devices were vacuumed for at least 20 minutes to remove all the air from the channels and traps. Syringes and falcon tubes were placed on the height control system and lines were connected as follows; media syringes were plugged first and kept above all other inputs to prevent media contamination. Adjusting the height of the cells-containing falcon tube as well as media and waste aids in controlling the cell seeding in the traps. Although many cells pass through the chip directly towards the waste port, few cells got captured via micro valves and seeded the traps. Once 10-20 cells were captured in each trapping region, the flow from the cell loading port was reverted by decreasing the height to the same level as for the auxiliary waste.

### Time-lapse imaging, growth conditions and data analysis

Live-cell imaging was performed on the Automated inverted Leica DMi8 fluorescence microscope equipped with Hamamatsu Orca Flash V3 camera that was controlled by μManager(micro-manager.org). Images were captured with 40x dry objective NA=0.8 (Leica Inc.).Traps containing cells were imaged every 10 minutes on three different channels (DIC, GFP (Excitation: 488 Emission: 515, and mCherry Excitation: 583, Emission: 610) with CoolLed pE600 LED excitation source and standard Chroma epifluorescent filter set. Experiments were run for up to 72 hours under the continuous supply of nutrients in the microfluidic device. Acquired images were initially processed in Fiji (https://imagej.net/Fiji) using custom scripting to extract positions with exponentially growing yeast cells. Constitutively expressed mCherry marker was used to identify healthy cells that are in the exponential growth phase.

Dead or non-growing individuals were discarded by correcting dEGFP signal according to the formula *dEGFP*/(*dEGFP* + *mCherry*). Each image was divided into 25 regions of interest (ROIs) and analyzed separately to isolate regions where cells were actively growing and could be tracked over time. The posterior analysis was done with custom R-studio scripts. Firstly, raw data were detrended using the detrend function from “pracma” R package and then smoothed with Savitzky-Golay Smoothing function (savgol), from the same package, with a filter length of 15 was applied and the signal was normalized between 0 and 1 to generate time-course plots and kymographs of traps. Amplitudes were calculated with find peaks within the Process Data using the “findpeaks” function from “pracma” R package with nups and ndowns of 6, and periods were calculated by calculating distances between dEGFP peaks. Precision of response was based on relative differences between peaks widths of stimulus and response, respectively. Precision per *i-th* trap was calculated as follows : *precision*^*i*^ = 1- (*width_output* ^*i*^ – *width_input*)/(*width_output*^*i*^ + *width_input*). Autocorrelations traces and power spectrum densities were calculated using standard calculations with Matlab derived packages autocorr and Fast Fourier Transformation (fft).

### Mathematical modeling

In the CrossTalker model, MarR (*M*) and IacR (*I*) levels as well as dEGFP reporter concentrations (R) were described using Delayed differential equations scheme (DDE) that was implemented in Matlab 2018b Inc. The dde23 DDE function was used to numerically solve for time-dependent concentrations of species. For simplicity, processes such as transcription, translation, and monomer oligomerization were captured in a single parameter (*α*_*o*_) whereas MarR degradation was modeled separately as basal(*δ*_*o*_) and phytohormone-dependent(*δ*_*sa*_ and *<u>δ</u>*_<u>*iaa*</u>_) rates:

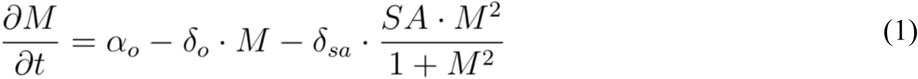

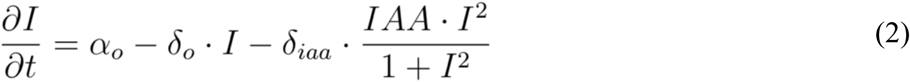

SA and IAA are normalized salicylic acid and auxin concentrations from 0(min) to 1(max) that follow stimulus pulses (forcing period *T*_*f*_) according to formulas:

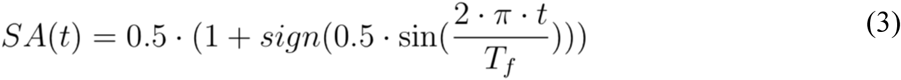

and,

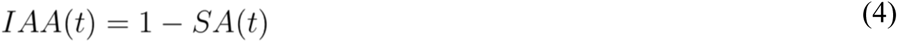

The time evolution of reporter gene dEGFP (*R*) was described as follows:

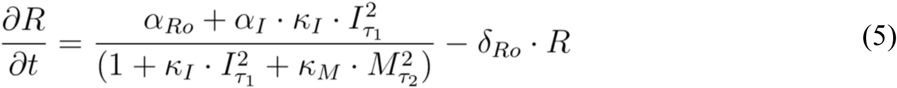

where *α*_*Ro*_ is a leakiness of the promoter, α_*I*_ is an IacR-dependent protein production, *κ*_*I*_ is the strength of IacR-dependent activation and *κ*_*M*_ is the strength of MarR-mediated repression. The parameter *δ*_*Ro*_ is dEGFP degradation rate. Model integrates two different delays for activation (*τ*_*1*_) and repression (*τ*_*2*_), respectively. Model parameters were scanned and predictions qualitatively compared to the experimental time-curse data (Fig. 2g). We found that only an extended delay in repression (τ2 >> τ1) closely mimic these experimental observations (Supplementary Fig. 5h). The same set of parameters was then used for generating Supplementary Fig. 5i with a 50% shorter period of stimulus (Supplementary Table 1).

## Supplementary Information

### Supplementary Figures 1-6 with legends

**Supplementary Figure 1.**
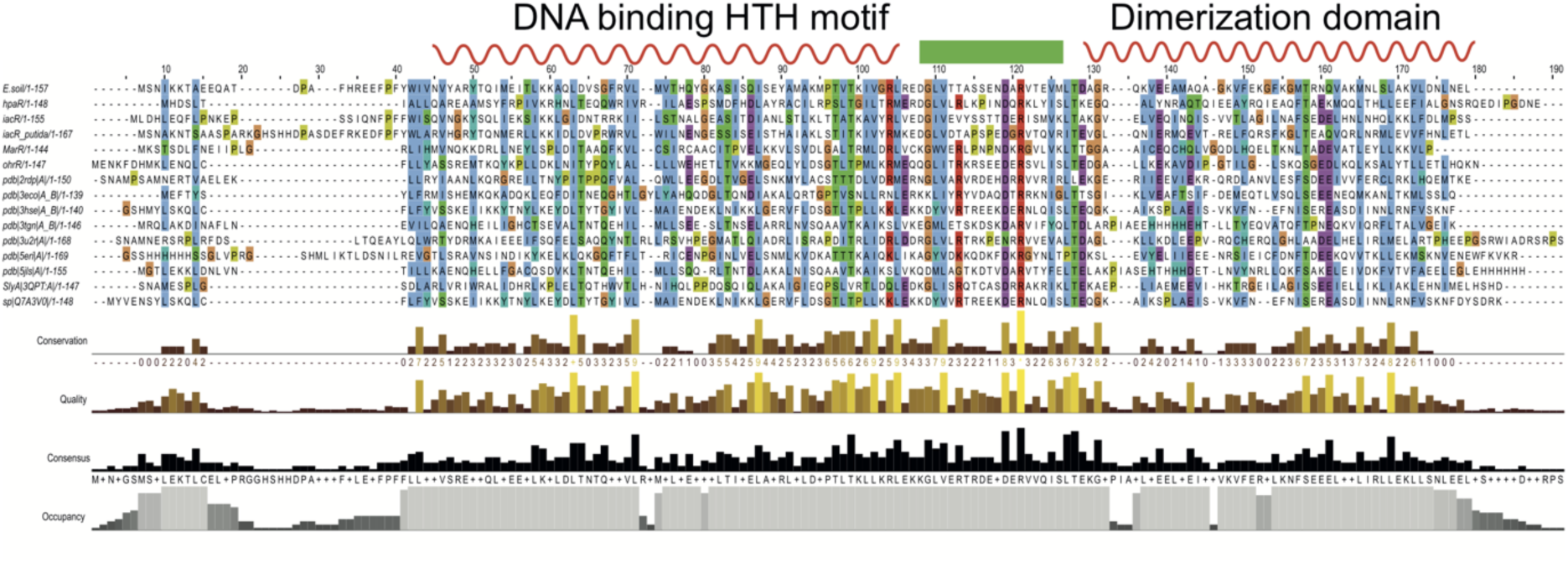
Common structural features of MarR-type repressors. Structural alignment of MarR-type transcriptional repressors highlights structurally conserved regions such as DNA binding and dimerization interfaces α-helix (red) : β-sheet (green) : α-helix (red).

**Supplementary Figure 2.**
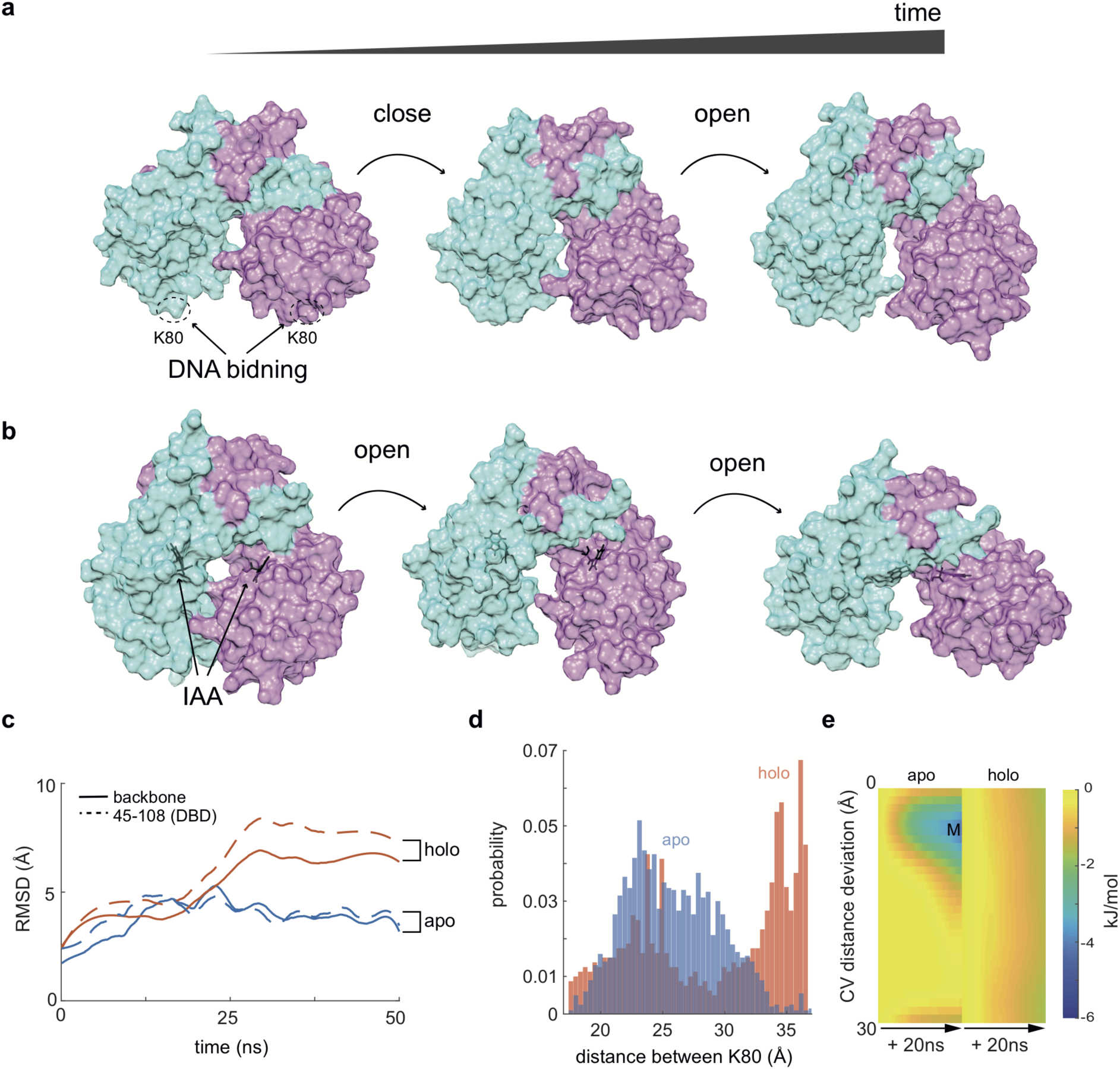
Predicted similarities between MarR and in the iacR auxin sensing mechanisms. (**a, b**), Time-lapse from MD simulations of the predicted iacR apoprotein (a) (monomers shown in cyan and purple) and IAA-bound holoprotein (b). DNA binding charged K80 residues (equivalent to R73 in MarR) are shown. Note cycles of closure and opening in the apo configuration similar to those observed in MarR simulations (Fig. 1a). Predicted dynamic poses of two IAA molecules along the time-course of MD simulation (b). Note, a progressive opening of the dimer (unlike MarR_holo_ which closes) which pushes pair of DNA binding domain far enough to preclude potential DNA binding. (**c**), RMSD of backbone, and putative DNA binding domain of iacR (residues 45-108) for apo and halo configuration are shown. (**d**), A probability distribution of the mean distance between K80 atoms of both monomers for apo (blue) and holo(red) iacR configurations. Note that holo state provides substantially longer distance between DNA binding domains compared to apo. (**e**), Energy landscapes for iacR configurations (apo, holo) for deviation from a putative distance of 20 A° between K80 (both chains) that could promote DNA binding. Heatmap of energy for apo and holo configurations indicate that unlike apoprotein, the IAA-bound holoprotein cannot reach the minimum required for association with the target DNA.

**\Supplementary Figure 3.**
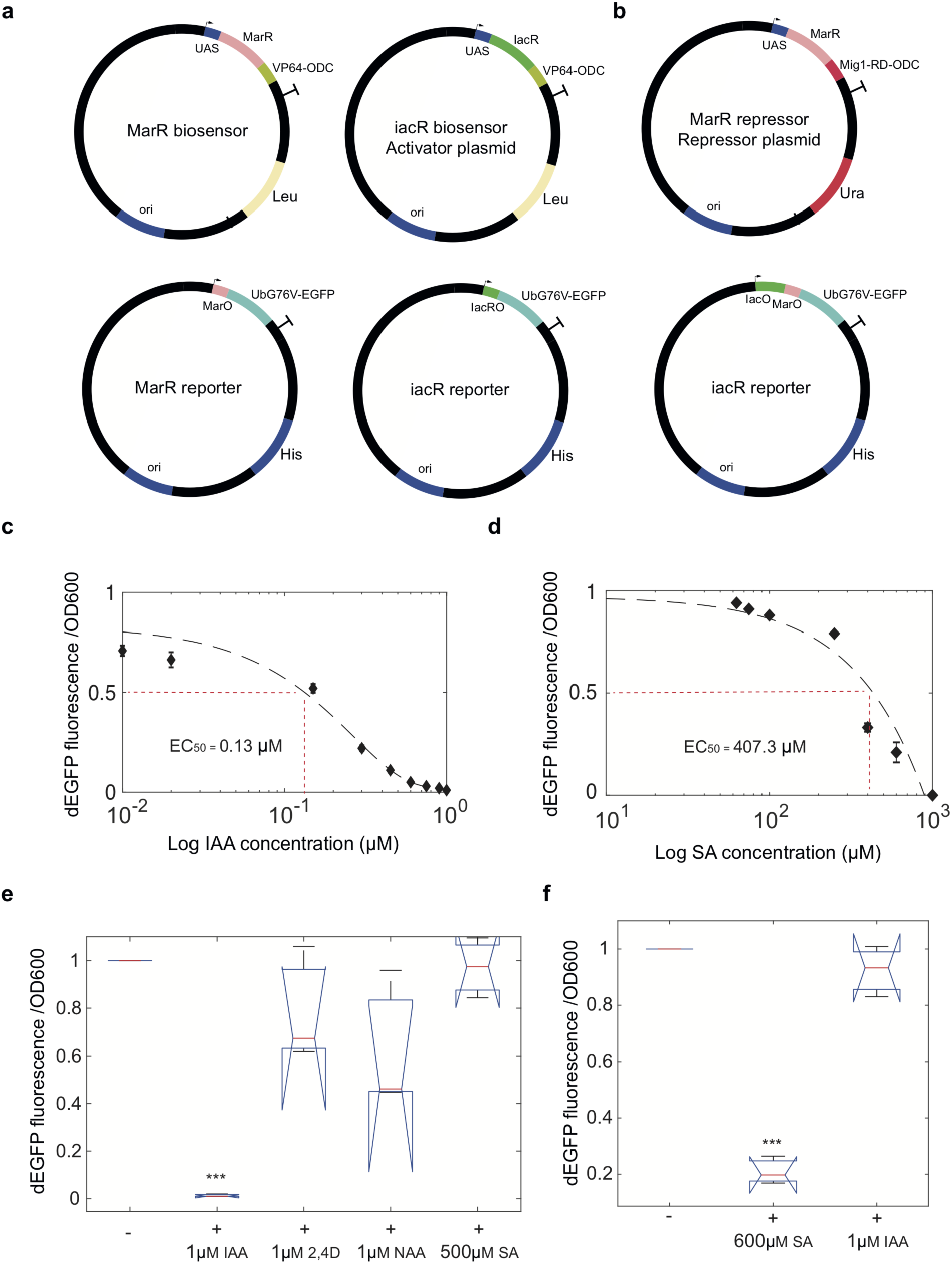
A description of experimental biosensing systems and their robust response in physiological ranges of phytohormone concentrations. (**a**), Plasmid compositions include all parts used to construct synthetic gene circuits and fluorescent reporters. UAS denotes GAL4 binding sites in Galactose inducible promoter. MarO and IacO are operator sequences for MarR and IacR transcriptional regulators. (**b**), Construction of MarR repressor and reporter plasmids are shown. MarR was tagged with the last 24 aa of MIG1 transcription factor that binds the general TUP1-SSN6 co-repressor complex in yeast. (**c, d**), Response curves of IacR module to IAA (c) and MarR module to SA (d) are shown. Note differences in a dynamic range (2-orders (c) and 1-order of magnitude (d)) towards physiological concentrations of phytohormones. Means and standard errors are from three independent experiments. (**e**), Orthogonality and high specificity of IacR module to IAA but not SA or other synthetic auxins (2,4D and NAA). (**f)**, MarR module is insensitive to IAA. A one-way ANOVA test was used to test the significance of three independent replicates (p-value *** < 0.001).

**Supplementary Figure 4.**
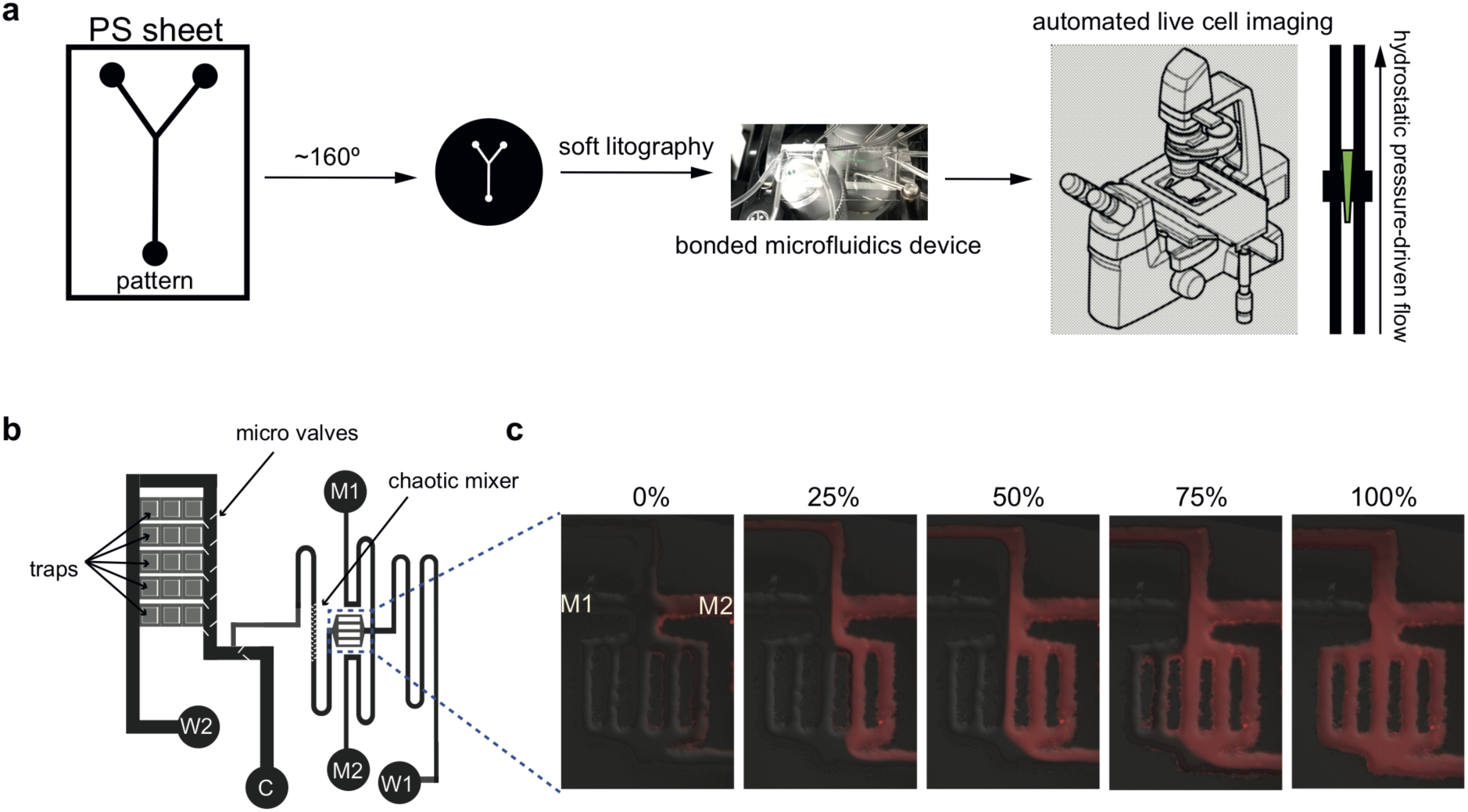
Microfluidics fabrication and flow control scheme. (**a**), A simplified process of microfluidic device fabrication. Initially, the pattern of channels is printed with a laser printer at 1200dpi resolution on plastic polystyrene (PS) sheet. Next, a PS sheet is baked at approx. 160°C such that it shrinks around 70% in x-y and increases in height by 10-fold. The height of channels and traps can be controlled by the density of ink deposition to create cell monolayers. Soft lithography is used to make PDMS silicon devices that are then bonded to the cover glass to finalize the device. Microfluidics platform integrates automated epifluorescent microscope and customized microfluidic flow system with linear actuators which allows the gravity-regulated hydrostatic pressure control. (**b**), A final design of microfluidic chip for experiments. Channels widths were 120μm for in the mixer module and 500μm in main channels. The approximate height of channels was 25 microns. Cell traps had 500μm x 500μm size and height of approximately 7 microns. M1 and M2 are media ports that supply media with or without phytohormones. W1 is a media waste port that controls an influx in the mixer module and W2 is a general waste port. Cells are loaded through C port and seeded in the trapping region with the help of microvalves. (**c**), Screenshots from mixing conditions and 3-step mixer calibration (0%, 50%, 100%) with Rhodamine B red fluorescent dye.

**Supplementary Figure 5.**
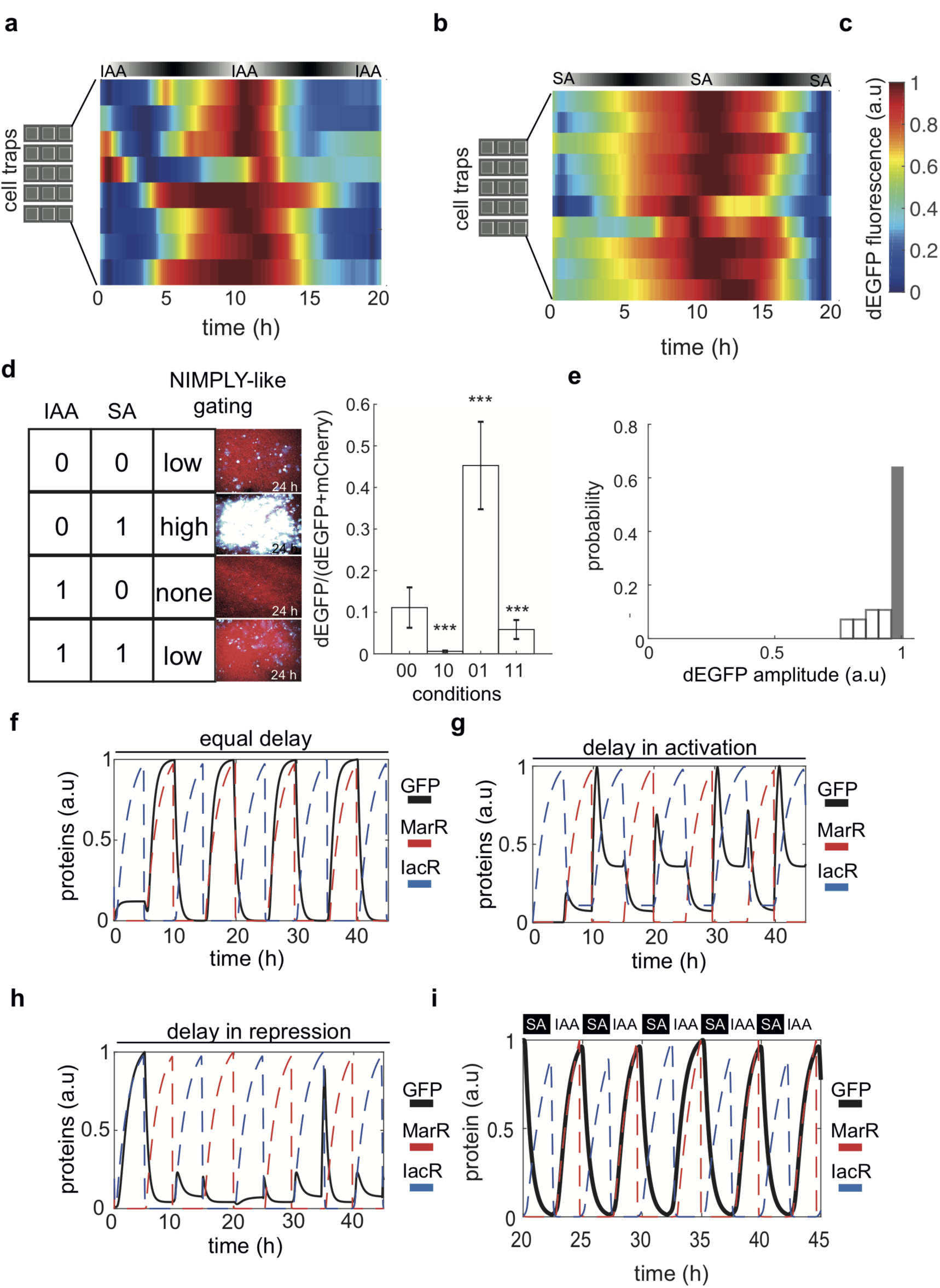
On-chip synchronization of single and crosstalk phytohormone sensing modules and simulations of Crosstalker computer model in different regimes. (**a, b**), Synchronization of single SA(a) or IAA(b) biosensing modules across trapping regions in microfluidics experiments related to Fig. 2b, c. (**c**) Color coding map for dEGFP fluorescence signal. (**d**), NIMPLY logic gate input table and its realization by CrossTalker circuit in microfluidics experiments. Examples of screenshots after 24 hours of growth in microfluidics traps n = 3 per condition are shown (ANOVA one-way p-value *** < 0.001). Standard errors (SE) are shown. dEGFP signal normalized to mCherry signal. (**e**), Histogram of the dEGFP fluorescence amplitude variation across all traps monitored on the microfluidic device, related to Fig. 2g. (**f-h**), Computer model simulations of CrossTalker circuit for three different scenarios: equal delay in activation and repression of dEGFP (f), longer delay in positive feedback only (g), and a longer delay in repression branch (h). The scenario presented in (h) is in the best agreement with experimental observation (Fig. 2g). (**i**), CrossTalker computer model simulations predict synchronized oscillations of a reporter gene in response to fast antiphase pulses of SA and IAA.

**Supplementary Figure 6.**
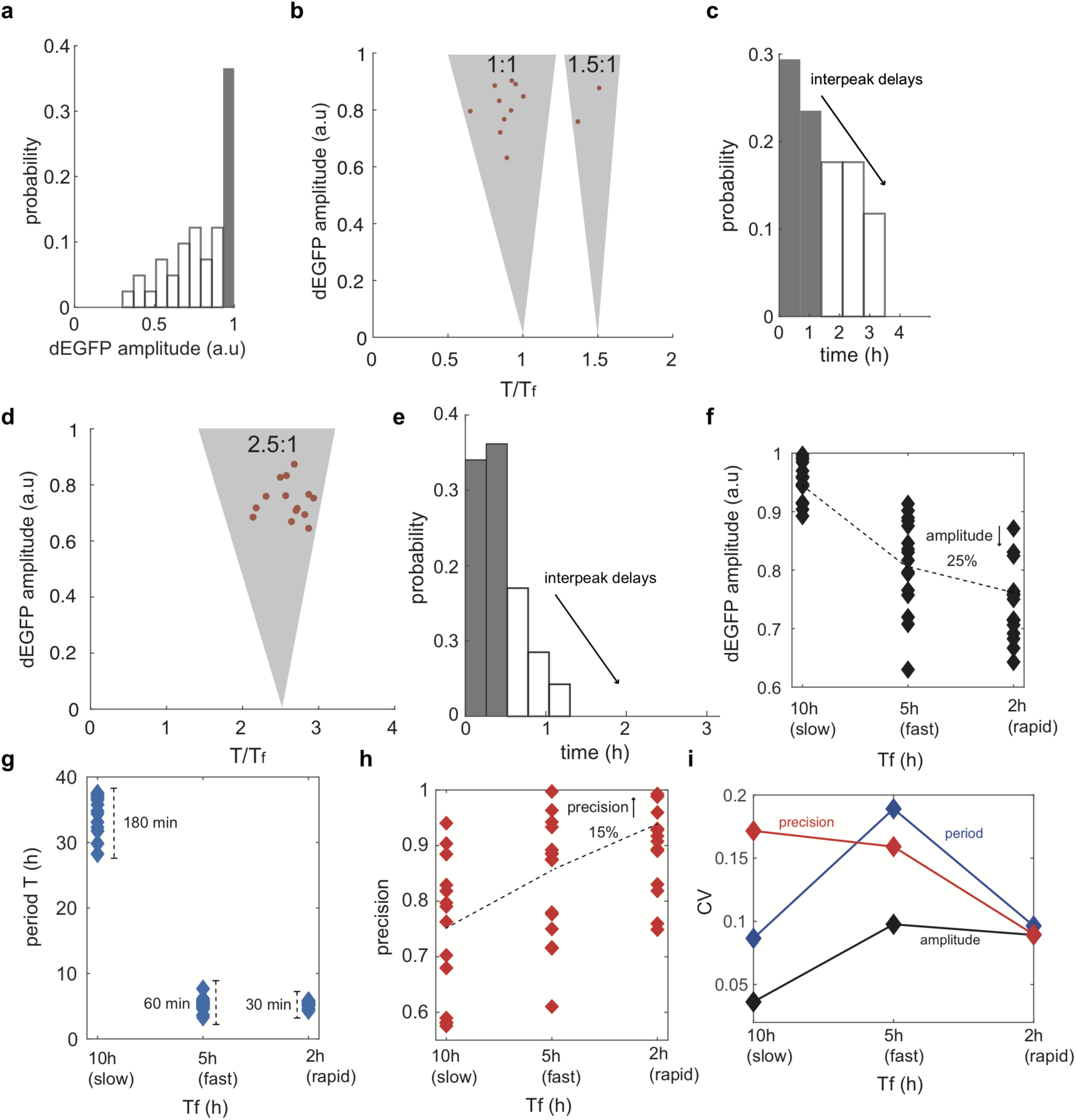
Dynamics of CrossTalker circuit in response to the increased frequency of hormone input. (**a**), Variation in the mean dEGFP amplitude across different traps as observed in Fig. 3b, c. (**b**), Synchronization regions of CrossTalker circuit in response to the 5h period of hormone stimulus. Two modes were observed in at least n=14 independent traps, dominant 1:1 and weak 1:1.5 locking between response periods (*T*) and forcing input period (*T*_*f*_). (**c**), Distribution of inter-peak delays for experiments presented in Fig. 3b, c demonstrates the phase diffusion that is correlated with increased exposure to phytohormones (‘capacitance’). (**d**), Synchronization regions of CrossTalker circuit in response to the 2h period of hormone stimulus. One dominant modes was observed: 2.5:1 locking between response (*T*) and forcing input periods (*T*_*f*_). (**e**), Distribution of inter-peak delays for experiments presented in Fig. 3f, g demonstrates a lack of significant delay and thus improved precision of the circuit response compared to (c). (**f)**, A reduction of response amplitude by 1/4 with an increased frequency of hormone input. (**g**), A consistency of period increased by a 2-fold with faster stimulus. (**h**), the response precision was improved by 15% with a higher frequency of hormone inputs. (**i**), The coefficient of variation (CV) for three response descriptors (amplitude in black, a period in blue and precision in red) under 3 different frequencies of hormone inputs. Note, the increased consistency of period and precision for the 2h cyclic hormone stimulus.

**Supplementary Table 1.** CrossTalker computer model parameters.

| Parameter | Value |
| --- | --- |
| $\alpha_o$ | 4 5 |
| $\delta_o$ | 0.5 |
| $\delta_{SA}$ | 10000 |
| $\delta_{IAA}$ | 100 |
| $\alpha_{Ro}$ | 0.01 |
| $\alpha_I$ | 10 10 |
| $k_I$ | 1 |
| $k_M$ | 10 |
| $\delta_{Ro}$ | 2 |
| $T_f$ | 10 ( Supplementary Fig. 5f-h); 5<br>( Supplementary Fig. 5i) 15 |
| $\tau_I$ | 5 |
| $\tau_2$ | 0.01 |

### Supplementary Movie legends

**Supplementary Movie 1**. Molecular Dynamics simulation of MarR apoprotein. The total time of the simulation was 50ns. Related to Fig. 1a.

**Supplementary Movie 2**. Molecular Dynamics simulation of MarR apoprotein associated with DNA operator site (black). The total time of the simulation was 50ns. Related to Fig. 1b.

**Supplementary Movie 3**. Molecular Dynamics simulation of MarR holoprotein associated with SA (black). The total time of the simulation was 50ns. Related to Fig. 1c.

**Supplementary Movie 4**. Microfluidic trap with cells carrying the CrossTalker circuit subjected to antithetic changes in both phytohormones with a 10-hour period. Related to Fig. 2g, h. Time changes of a hormone stimulus are shown on the panel above the movie (black(max SA, min IAA), white (max IAA, min SA))as shown in Fig. 2g.

**Supplementary Movie 5**. Increased responsiveness of the CrossTalker circuit subjected to antithetic pulses of both phytohormones with 5 hour period (two traps are shown simultaneously). Related to Fig. 3b, c. Time changes of a hormone stimulus are shown on the panel above the movie (black(max SA, min IAA), white (max IAA, min SA)) as shown in Fig. 3b.

**Supplementary Movie 6**. Increased responsiveness of the CrossTalker circuit subjected to antithetic pulses of both phytohormones with 2 hour period. Related to Fig. 3f, g. Time changes of a hormone stimulus are shown on the panel above the movie (black(max SA, min IAA), white (max IAA, min SA)) as shown in Fig. 3f.

## References

1. Robert-Seilaniantz, A., Grant, M., & Jones, J. D. G. (2011). Hormone Crosstalk in Plant Disease and Defense: More Than Just JASMONATE-SALICYLATE Antagonism. Annual Review of Phytopathology, 49(1), 317–343.

2. Santner, A., Calderon-Villalobos, L. I. A., & Estelle, M. (2009). Plant hormones are versatile chemical regulators of plant growth. Nature Chemical Biology, 5(5), 301–307.

3. Blázquez, M. A., Nelson, D. C., & Weijers, D. (2020). Evolution of Plant Hormone Response Pathways. Annual Review of Plant Biology, 71(1), 327–353.

4. Brunoud, G., Wells, D. M., Oliva, M., Larrieu, A., Mirabet, V., Burrow, A. H., Beeckman, T., Kepinski, S., Traas, J., Bennett, M. J., & Vernoux, T. (2012). A novel sensor to map auxin response and distribution at high spatio-temporal resolution. Nature, 482(7383), 103–106.

5. Liao, C.-Y., Smet, W., Brunoud, G., Yoshida, S., Vernoux, T., & Weijers, D. (2015). Reporters for sensitive and quantitative measurement of auxin response. Nature Methods, 12(3), 207–210.

6. Huang, W. E., Huang, L., Preston, G. M., Naylor, M., Carr, J. P., Li, Y., Singer, A. C., Whiteley, A. S., & Wang, H. (2006). Quantitative in situ assay of salicylic acid in tobacco leaves using a genetically modified biosensor strain of Acinetobacter sp. ADP1. The Plant Journal, 46(6), 1073–1083.

8. Pierre-Jerome, E., Jang, S. S., Havens, K. A., Nemhauser, J. L., & Klavins, E. (2014). Recapitulation of the forward nuclear auxin response pathway in yeast. PNAS, 111(26), 9407–9412.

9. Parízková, B., Pernisová, M., & Novák, O. (2017). What Has Been Seen Cannot Be Unseen—Detecting Auxin In Vivo. International Journal of Molecular Sciences, 18(12), 2736.

10. Uchida, N., Takahashi, K., Iwasaki, R., Yamada, R., Yoshimura, M., Endo, T. A., Kimura, S., Zhang, H., Nomoto, M., Tada, Y., Kinoshita, T., Itami, K., Hagihara, S., & Torii, K. U. (2018). Chemical hijacking of auxin signaling with an engineered auxin–TIR1 pair. Nature Chemical Biology, 14(3), 299–305.

11. Li, S., Prasanna, X., Salo, V. T., Vattulainen, I., & Ikonen, E. (2019). An efficient auxin-inducible degron system with low basal degradation in human cells. Nature Methods, 16(9), 866–869.

12. Deochand, D. K., & Grove, A. (2017). MarR family transcription factors: dynamic variations on a common scaffold. Critical Reviews in Biochemistry and Molecular Biology, 52(6), 595–613.

13. Alekshun, M. N., Levy, S. B., Mealy, T. R., Seaton, B. A., & Head, J. F. (2001). The crystal structure of MarR, a regulator of multiple antibiotic resistance, at 2.3 A resolution. Nature Structural Biology, 8(8), 710–714.

14. Hao, Z., Lou, H., Zhu, R., Zhu, J., Zhang, D., Zhao, B. S., Zeng, S., Chen, X., Chan, J., He, C., & Chen, P. R. (2014). The multiple antibiotic resistance regulator MarR is a copper sensor in Escherichia coli. Nature Chemical Biology, 10(1), 21–28.

15. Shu, H.-Y., Lin, L.-C., Lin, T.-K., Chen, H.-P., Yang, H.-H., Peng, K.-C., & Lin, G.-H. (2015). Transcriptional regulation of the iac locus from Acinetobacter baumannii by the phytohormone indole-3-acetic acid. Antonie van Leeuwenhoek, 107(5), 1237–1247.

16. Seipel, K., Georgiev, O., & Schaffner, W. (1992). Different activation domains stimulate transcription from remote (‘enhancer’) and proximal (‘promoter’) positions. The EMBO Journal, 11(13), 4961–4968.

17. Takeuchi, J., Chen, H., Hoyt, M. A., & Coffino, P. (2008). Structural elements of the ubiquitin-independent proteasome degron of ornithine decarboxylase. Biochemical Journal, 410(2), 401–407.

18. Stack, J. H., Whitney, M., Rodems, S. M., & Pollok, B. A. (2000). A ubiquitin-based tagging system for controlled modulation of protein stability. Nature Biotechnology, 18(12), 1298–1302.

19. Grimes, A., Breslauer, D. N., Long, M., Pegan, J., Lee, L. P., & Khine, M. (2008). Shrinky-Dink microfluidics: rapid generation of deep and rounded patterns. Lab on a Chip, 8(1), 170–172.

20. Ostling, J., Carlberg, M., & Ronne, H. (1996). Functional domains in the Mig1 repressor. Molecular and Cellular Biology, 16(3), 753 LP–761.

21. Lavy, M., & Estelle, M. (2016). Mechanisms of auxin signaling. Development (Cambridge, England), 143(18), 3226–3229.

22. Weijers, D., & Wagner, D. (2016). Transcriptional Responses to the Auxin Hormone. Annual Review of Plant Biology, 67, 539–574.

23. Kumar, D. (2014). Salicylic acid signaling in disease resistance. Plant Science : An International Journal of Experimental Plant Biology, 228, 127–134.

24. Vlot, A. C., Dempsey, D. A., & Klessig, D. F. (2009). Salicylic Acid, a multifaceted hormone to combat disease. Annual Review of Phytopathology, 47, 177–206.

25. Tan, S., Abas, M., Verstraeten, I., Glanc, M., Molnár, G., Hajný, J., Lasák, P., Petrík, I., Russinova, E., Petrášek, J., Novak, O., Pospisil, J., & Friml, J. (2020). Salicylic Acid Targets Protein Phosphatase 2A to Attenuate Growth in Plants. Current Biology, 30.

26. Pasternak, T., Groot, E. P., Kazantsev, F. V, Teale, W., Omelyanchuk, N., Kovrizhnykh, V., Palme, K., & Mironova, V. V. (2019). Salicylic Acid Affects Root Meristem Patterning via Auxin Distribution in a Concentration-Dependent Manner. Plant Physiology, 180(3), 1725 LP–1739.

27. Gibson, D. G., Young, L., Chuang, R.-Y., Venter, J. C., Hutchison, C. A., & Smith, H. O. (2009). Enzymatic assembly of DNA molecules up to several hundred kilobases. Nature Methods, 6(5), 343–345.

